# Extensive disruption of HoxA/D genes does not cause obvious defects in anterior vertebral patterning in zebrafish

**DOI:** 10.64898/2026.09.09.750522

**Authors:** Taisei Tani, Akiteru Maeno, Hidemichi Nakazawa, Towa Fujibayashi, Kaito Suda, Akinori Kawamura

## Abstract

Hox genes play central roles in specifying positional identities along the vertebrate anterior-posterior axis. In mice, genetic analyses have demonstrated that Hox genes distributed among the four Hox clusters contribute to vertebral patterning, with extensive functional redundancy among paralogous genes. Our previous genetic analysis in zebrafish identified important roles for HoxB- and HoxC-related genes in specifying anterior vertebral identities, whereas the contributions of HoxA- and HoxD-related genes remained unresolved. Here, we examined adult zebrafish carrying extensive combinations of *hoxaa, hoxab*, and *hoxda* cluster deletions and generated five-gene homozygous mutants carrying frameshift mutations in *hoxa3a, hoxa4a, hoxa5a, hoxd3a*, and *hoxd4a*. X-ray micro-CT analysis revealed no obvious anterior vertebral transformations in either the compound cluster mutants or the five-gene mutants. Together with our previous findings, these results suggest that vertebral patterning functions are distributed unevenly among zebrafish hox clusters, with a predominant contribution from HoxB/C-related genes.

## Text

Hox genes encode homeodomain transcription factors that specify positional identities along the anterior-posterior axis of vertebrate embryos. Genetic studies in mice have established that Hox genes belonging to multiple paralog groups and distributed among the four Hox clusters contribute to vertebral patterning (Wellik, 2007). Loss of *Hoxd3* alters the identities of the first and second cervical vertebrae, including the atlas and axis, whereas *Hoxa4* mutants show transformations within the cervical region (Condie and Capecchi, 1993; Horan et al., 1994). Importantly, compound mutations of paralogous genes located in different Hox clusters produce more extensive and dose-dependent transformations. For example, combined disruption of Hox3 or Hox4 paralogs markedly enhances cervical vertebral phenotypes, demonstrating substantial functional redundancy among Hox genes residing in different clusters (Condie and Capecchi, 1994; Horan et al., 1995). Thus, in the mouse, vertebral identities are specified by overlapping activities of Hox genes distributed among different clusters rather than by the action of a single Hox cluster.

Teleost fishes underwent an additional whole-genome duplication, and zebrafish retain seven hox clusters (Amores et al., 1998). Zebrafish hox genes exhibit extensive, nested expression domains along the body axis (Prince et al., 1998), but axial expression does not necessarily establish a functional requirement for a corresponding vertebral identity. In our previous loss-of-function study, several HoxB- and HoxC-related genes were shown to contribute to anterior vertebral patterning in zebrafish (Maeno et al., 2024). Mutations in these genes produced distinct changes in anterior vertebral morphology, providing direct genetic evidence that a subset of the zebrafish hox repertoire is functionally required for vertebral identity. Importantly, the affected vertebral elements differed among the HoxB/C mutants, indicating that these phenotypes do not simply reflect a nonspecific skeletal defect but instead reveal gene-specific contributions to anterior vertebral patterning. This previous analysis therefore provided a functional framework for asking whether HoxA- and HoxD-related genes make comparable contributions. Zebrafish retain two HoxA-related clusters, *hoxaa* and *hoxab*, but only one HoxD-related cluster, *hoxda* (Amores et al., 1998). Previous whole-cluster deletion analyses did not reveal obvious anterior vertebral abnormalities in *hoxaa* or *hoxda* homozygous mutants (Yamada et al., 2021), but these experiments could not exclude functional redundancy among HoxA/D-related genes. Moreover, more extensive combinations of HoxA/D-related cluster deletions do not survive to adulthood (Ishizaka et al., 2024). We therefore examined the strongest viable combinations of HoxA/D-related cluster deletions and, in parallel, generated compound mutations in five HoxA/D genes belonging to paralog groups relevant to the anterior vertebral region.

Zebrafish are otophysan fishes in which the anterior vertebrae are associated with specialized skeletal elements that form the Weberian apparatus. In particular, the first four vertebrae and their associated Weberian ossicles have distinct morphologies, providing readily identifiable anatomical landmarks for assessing individual anterior vertebral identities (Bird and Mabee, 2003; Grande and Young, 2004; Maeno et al., 2024). The *hoxaa, hoxab*, and *hoxda* cluster-deficient alleles were generated previously (Yamada et al., 2021). Because fish homozygous for more extensive combinations of these HoxA/D-related cluster deletions do not survive to adulthood, we analyzed the viable adult genotypes *hoxaa*^*−/−*^;*hoxab*^*+/−*^;*hoxda*^*+/−*^ and *hoxaa*^*+/−*^;*hoxab*^*+/−*^;*hoxda*^*−/−*^ (Ishizaka et al., 2024). Pelvic fin abnormalities, including unilateral or bilateral fin loss, have previously been reported in these compound cluster mutant genotypes (Nakazawa et al., 2026). In contrast, X-ray micro-CT analysis revealed no obvious alterations in the morphology or identity of the anterior vertebrae or associated Weberian apparatus in any of the specimens examined (*n* = 3 per mutant genotype; Fig. 1; Supplementary Movies 1–3). Thus, despite their effects on pelvic fin development, these extensive HoxA/D-related cluster deletions did not cause obvious anterior vertebral transformations in the adult mutants examined.

**Figure 1.**
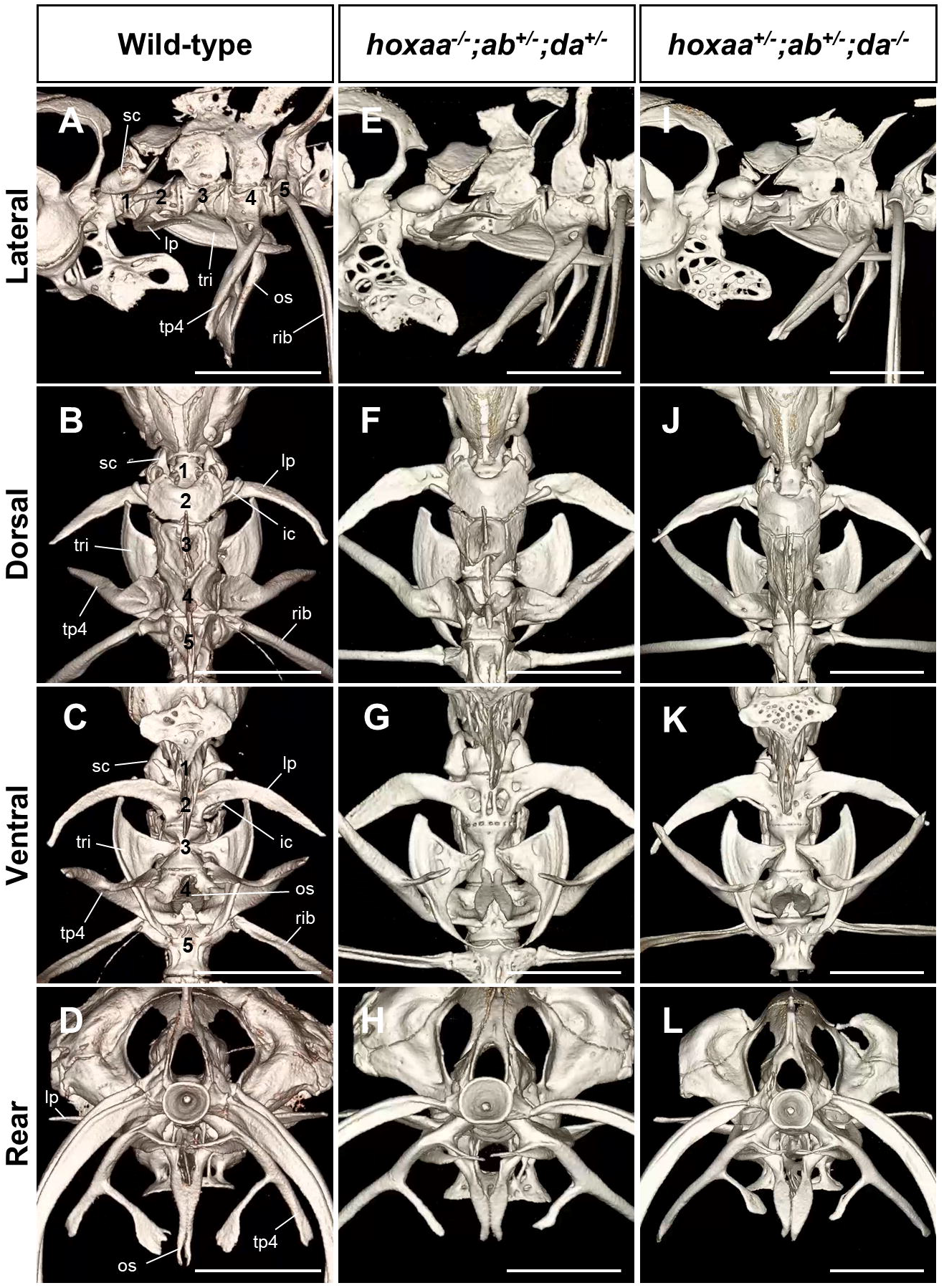
Anterior vertebral morphology is preserved in compound HoxA/D-related cluster mutants. X-ray micro-CT images of the anterior vertebral column of wild-type, *hoxaa*^−/−^;*hoxab*^+/−^;*hoxda*^+/−^, and *hoxaa*^+/−^;*hoxab*^+/−^;*hoxda*^−/−^ adult zebrafish. Characteristic skeletal elements associated with the first four vertebrae are indicated in the wild-type panels: scaphium (sc), lateral process (lp), intercalarium (ic), tripus (tri), os suspensorium (os), and transverse process of vertebra 4 (tp4). Three fish were analyzed for each mutant genotype, and no obvious alterations in the morphology or identity of the anterior vertebrae or Weberian apparatus were detected (*n* = 3 per genotype). Three-dimensional reconstructions of the anterior vertebral column of wild-type, *hoxaa*^−/−^;*hoxab*^+/−^;*hoxda*^+/−^, and *hoxaa*^+/−^;*hoxab*^+/−^;*hoxda*^−/−^ fish are shown in Supplementary Movies 1, 2, and 3, respectively. Scale bar: 1 mm.

Because these viable compound cluster mutants retain functional HoxA/D alleles, the absence of an obvious phenotype could still reflect redundancy among the remaining genes. The zebrafish *hoxab* cluster lacks paralogs 3–8 (Fig. 2A; Amores et al., 1998; Yamada et al., 2021). We therefore focused on five HoxA/D genes belonging to the relevant anterior paralog groups: *hoxa3a, hoxa4a, hoxa5a, hoxd3a*, and *hoxd4a*. CRISPR-Cas9 targeting generated a compound *hoxa3a;a4a;a5a* allele carrying frameshift mutations in all three genes on the same chromosome and a compound *hoxd3a;d4a* allele carrying frameshift mutations in both genes on the same chromosome (Fig. 2B, C). Crossing these two lines allowed us to obtain fish homozygous for both compound alleles, hereafter referred to as *hoxa3a;a4a;a5a;d3a;d4a* mutants. Unlike the more extensive HoxA/D cluster deletion combinations, these five-gene mutants were viable and survived to adulthood. Six adult homozygous mutants were analyzed by X-ray micro-CT. No obvious alterations in the morphology or identity of the anterior vertebrae, including the skeletal elements of the Weberian apparatus, were detected in any of the six specimens (*n* = 6; Fig. 3; Supplementary Fig. S1; Supplementary Movie 4). Thus, the major anterior vertebral landmarks were preserved in all six adult five-gene mutants examined.

**Figure 2.**
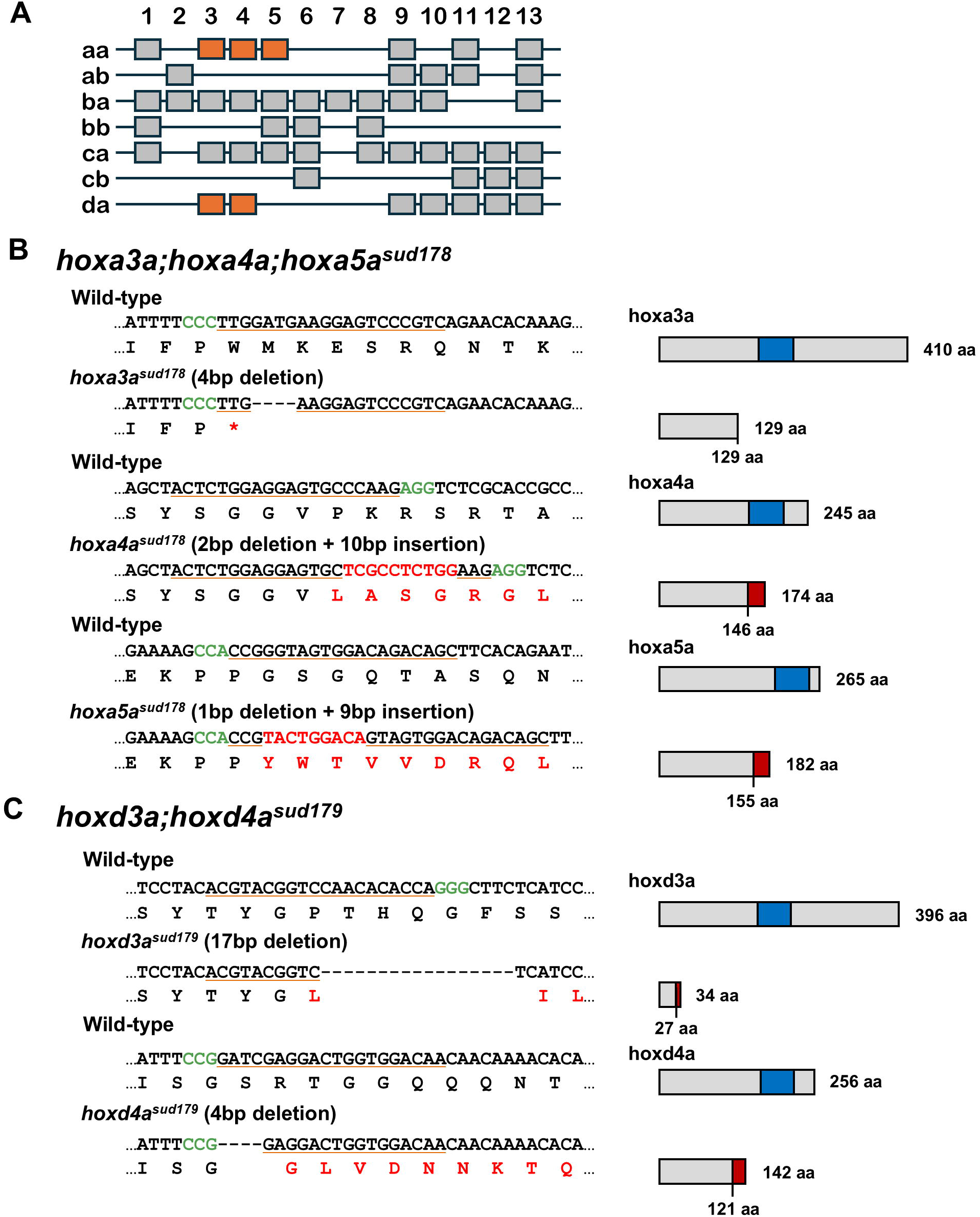
Generation of *hoxa3a;a4a;a5a* and *hoxd3a;d4a* compound mutant alleles. (A) Schematic representation of the zebrafish hox clusters showing the positions of *hoxa3a, hoxa4a, hoxa5a, hoxd3a*, and *hoxd4a* targeted in this study. Targeted genes are highlighted in orange. (B) CRISPR-Cas9-induced mutations in *hoxa3a, hoxa4a*, and *hoxa5a*. Wild-type and mutant nucleotide sequences surrounding each target site and the corresponding predicted amino acid sequences are shown. A compound *hoxa3a;a4a;a5a* allele carrying frameshift mutations in all three genes on the same chromosome was obtained. (C) CRISPR-Cas9-induced mutations in *hoxd3a* and *hoxd4a*. Wild-type and mutant nucleotide sequences surrounding each target site and the corresponding predicted amino acid sequences are shown. A compound *hoxd3a;d4a* allele carrying frameshift mutations in both genes on the same chromosome was obtained. PAM sequences are shown in green; altered nucleotide and predicted amino acid sequences resulting from the mutations are shown in red. Predicted protein structures are shown on the right, with homeodomains indicated in blue.

**Figure 3.**
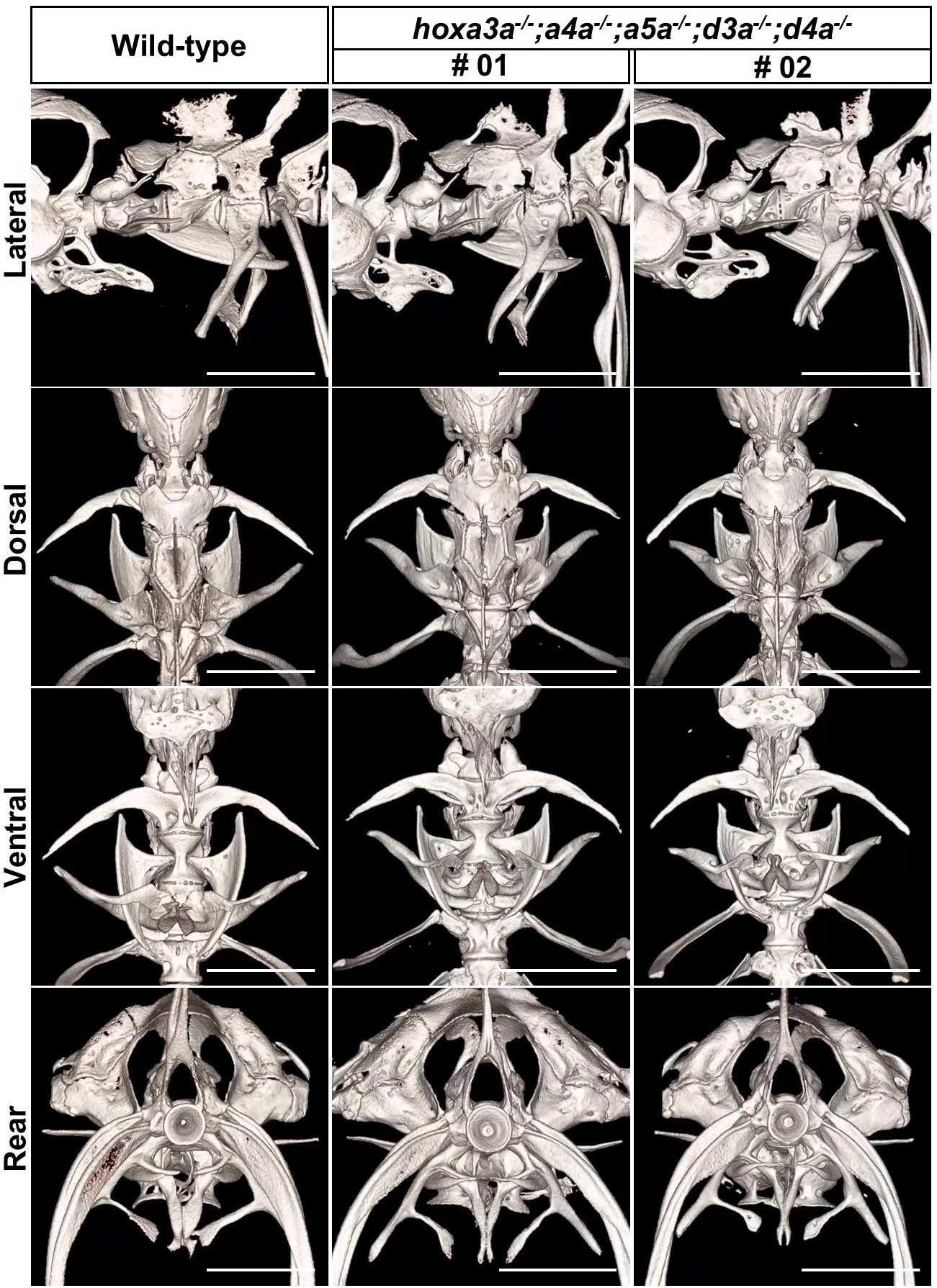
Anterior vertebral morphology in *hoxa3a;a4a;a5a;d3a;d4a* homozygous mutants. X-ray micro-CT reconstructions of the anterior vertebral column and associated Weberian apparatus of wild-type zebrafish and two individual adult *hoxa3a;a4a;a5a;d3a;d4a* homozygous mutants (#1 and #2). Six homozygous mutants were examined in total (*n* = 6), and no obvious alterations in anterior vertebral morphology or Weberian ossicle identity were detected in any of the specimens. A three-dimensional reconstruction of the anterior vertebral column of mutant #1 is shown in Supplementary Movie 4. CT reconstructions of the remaining four mutants (#3–6) are presented in Supplementary Fig. S1. Scale bar: 1 mm.

The preservation of anterior vertebral morphology in the five-gene mutants is particularly informative when compared with the pronounced abnormalities caused by disruption of HoxB/C-related genes. In our previous study, micro-CT analysis readily revealed transformations in the morphology and identity of individual anterior vertebrae and their associated Weberian elements (Maeno et al., 2024). Using a comparable approach, we detected no obvious anterior vertebral transformations in any of the six adult *hoxa3a;a4a;a5a;d3a;d4a* mutants examined. This contrast indicates that the HoxA/D genes investigated here do not make the same major contribution to anterior vertebral identity as the HoxB/C-related genes identified in our earlier study.

Together, these findings indicate that the genetic requirements for anterior vertebral patterning are distributed unevenly among zebrafish hox clusters, with predominant contributions from HoxB/C-related genes. This differs from the situation in mice, in which Hox genes from multiple clusters contribute to cervical vertebral identity through extensive cross-cluster redundancy. Loss of *Hoxd3* alters the atlas and axis, loss of *Hoxa4* affects cervical vertebral identities, and combined disruption of Hox3 or Hox4 paralogs produces stronger, dose-dependent transformations (Condie and Capecchi, 1993, 1994; Horan et al., 1994, 1995).

The zebrafish findings also fit a broader pattern of functional specialization among hox clusters. Whole-cluster mutant analyses revealed marked functional differences among the seven zebrafish hox clusters (Yamada et al., 2021). Subsequent studies demonstrated predominant roles of HoxC-related genes in dorsal and anal fin patterning (Adachi et al., 2024), essential roles of HoxB-derived clusters in pectoral fin formation (Kikuchi et al., 2025) and posterior pharyngeal pouch development (Toyama et al., 2026), and important contributions of HoxA/D-related clusters to pectoral fin growth (Ishizaka et al., 2024). More recently, HoxA/D paralog group 9–10 genes were shown to be essential for pelvic fin initiation (Nakazawa et al., 2026). Thus, the absence of obvious anterior vertebral transformations in HoxA/D mutants does not indicate a general lack of HoxA/D function. Rather, these findings demonstrate that distinct developmental functions are concentrated in different hox clusters in zebrafish.

Our findings also underscore the distinction between Hox gene expression and genetically demonstrated function. Numerous zebrafish hox genes exhibit nested expression domains along the anterior–posterior axis (Prince et al., 1998), yet these expression patterns do not necessarily predict which genes are required for the morphological identity of individual vertebrae. Comparative studies offer an instructive perspective. In the skate *Leucoraja erinacea*, boundaries of several Hox expression domains correlate with transitions between morphologically distinct vertebral regions (Criswell et al., 2021). By contrast, Oulion et al. (2011) described nested anterior expression boundaries of *HOX3, HOX4*, and *HOX5* within the morphologically similar anterior monospondylous vertebrae of the catshark *Scyliorhinus canicula*. These observations suggest that axial Hox regionalization may have preceded the emergence of conspicuous vertebral regionalization. Our genetic results extend this distinction: an axial Hox expression domain does not, by itself, establish that the corresponding gene is required for the formation of a particular vertebral identity. A functional vertebral Hox code must therefore be defined through genetic analysis, rather than inferred from expression boundaries alone.

The difference between zebrafish and mice raises an evolutionary question: when and how did Hox genes acquire their present contributions to vertebral identity? In mice, HoxA/D genes contribute to the specification of cervical vertebral identities, a region that underwent substantial morphological and functional elaboration during tetrapod evolution. One possibility is that vertebral patterning functions were initially concentrated in a subset of Hox clusters and that the contributions of HoxA/D genes expanded during tetrapod evolution, accompanying the elaboration of the cervical vertebral region. Such an expansion would not necessarily require the acquisition of new Hox expression domains. Instead, changes in the responses of downstream skeletal patterning networks to pre-existing Hox inputs could have increased the functional contributions of HoxA/D genes to vertebral identity (Woltering and Duboule, 2015). Alternatively, broader cross-cluster contributions may have been ancestral and subsequently reduced in the teleost lineage. Distinguishing these scenarios will require comparative genetic analyses across vertebrate lineages.

Several limitations should be considered. The viable compound cluster mutants retain some functional HoxA/D alleles, and residual activity or genetic compensation in the five-gene mutants cannot be completely excluded (Rossi and Kontarakis, 2022). Furthermore, our analyses were restricted to surviving adults, as more extensive HoxA/D cluster deletions are lethal before adulthood (Ishizaka et al., 2024). Nevertheless, all six adult five-gene mutants retained the major anterior vertebral landmarks, in striking contrast to the transformations observed after disruption of HoxB/C-related genes. Together, these findings demonstrate that the genetic requirements for anterior vertebral patterning are concentrated predominantly in HoxB/C-related genes in zebrafish, revealing a marked difference from the extensive cross-cluster functional redundancy established in mice.

## Methods

### Zebrafish

RIKEN wild-type (RW) zebrafish were obtained from the National BioResource Project Zebrafish (NBRP Zebrafish) and maintained at 27°C under a 14 h light/10 h dark cycle. Embryos were obtained by natural spawning. The *hoxaa*^*sud111*^, *hoxab*^*sud112*^, and *hoxda*^*sud116*^ cluster-deficient lines were generated previously (Yamada et al., 2021). Compound mutants carrying different combinations of these cluster deletions were generated by intercrossing the mutant lines as described previously (Ishizaka et al., 2024). All procedures involving live zebrafish were approved by the Animal Care and Use Committee of Saitama University and were conducted in accordance with institutional guidelines.

### Generation of *hoxa3a;a4a;a5a;d3a;d4a* homozygous mutants

Frameshift mutations in *hoxa3a, hoxa4a, hoxa5a, hoxd3a*, and *hoxd4a* were newly generated in this study using CRISPR–Cas9. To generate *hoxa3a;a4a;a5a* compound mutants, three guide RNAs targeting *hoxa3a, hoxa4a*, and *hoxa5a* were simultaneously introduced into one-cell-stage embryos. Germline screening identified a mutant carrying frameshift mutations in all three genes on the same chromosome (*sud178*). Similarly, two guide RNAs targeting *hoxd3a* and *hoxd4a* were co-introduced into one-cell-stage embryos. Germline screening identified a *hoxd3a;d4a* mutant carrying frameshift mutations in both genes on the same chromosome (*sud179*). The positions and sequences of these mutations are shown in Fig. 2. The crRNA sequences used for genome editing are listed in Supplementary Table 1. Fish carrying the *hoxa3a;a4a;a5a* and *hoxd3a;d4a* mutations were crossed, and their progeny were subsequently intercrossed to generate *hoxa3a;a4a;a5a;d3a;d4a* homozygous mutants. The *hoxa3a;a4a;a5a* and *hoxd3a;d4a* mutant lines have been cryopreserved as sperm at the National BioResource Project Zebrafish (NBRP Zebrafish) and are available upon request.

### Genotyping

Genomic DNA was prepared from caudal fin clips of anesthetized juvenile or adult zebrafish using the NaOH method. The *hoxaa, hoxab*, and *hoxda* cluster-deficient alleles were genotyped by PCR as described previously (Ishizaka et al., 2024; Yamada et al., 2021). The *hoxa3a;a4a;a5a* and *hoxd3a;d4a* mutations were identified by PCR amplification of the targeted regions followed by DNA sequencing. Primer sequences used for genotyping are listed in Supplementary Table 2. Genotypes of all fish used for micro-CT analysis were confirmed by DNA sequencing.

### X-ray micro-CT analysis

The anterior vertebral column of adult zebrafish was analyzed by X-ray micro-computed tomography essentially as described previously (Maeno et al., 2024). Adult fish were fixed in 4% paraformaldehyde in PBS overnight and transferred to 70% ethanol. Specimens were scanned using an X-ray micro-CT system (ScanXmate-E090S105; manufactured by Comscantechno Co., Ltd., Yokohama, Japan; currently maintained and technically supported by Voxel Works Co., Ltd., Tokyo, Japan) at a tube voltage of 85 kV and a tube current of 90 μA. Specimens were rotated through 360° in 0.24° steps, generating 1500 projection images. CT datasets were reconstructed using coneCTexpress software (Comscantechno) at an isotropic resolution of 5.0 μm. Three-dimensional images of the anterior vertebrae and Weberian apparatus were examined using OsiriX MD (Pixmeo). A total of six fully mature adult *hoxa3a;a4a;a5a;d3a;d4a* homozygous mutants were analyzed. The anterior vertebrae and Weberian ossicles were examined for morphological abnormalities and changes in skeletal identity.

## Acknowledgments

We thank the National BioResource Project (NBRP) Zebrafish for providing the RW strain and preserving the hox mutant lines. This work was supported by a Grant-in-Aid for Scientific Research (KAKENHI) from the Japan Society for the Promotion of Science (JSPS) (23K05790 to A.K.) and the National Institute of Genetics under the Joint Research Program (NIG-JOINT; 66A2021, 18A2022, 31A2023, 26A2024, 3B2025 to A.K.).

## Data Availability Statement

The *hoxa3a;a4a;a5a*^*sud178*^ and *hoxd3a;d4a*^*sud179*^ mutant lines generated in this study have been cryopreserved as sperm at NBRP Zebrafish and are available upon request (https://shigen.nig.ac.jp/zebrafish/). All other data supporting the findings of this study are available within the article and its Supporting Information.

