## Supplemental data for "Extensive disruption of HoxA/D genes does not cause obvious defects in anterior vertebral patterning in zebrafish"

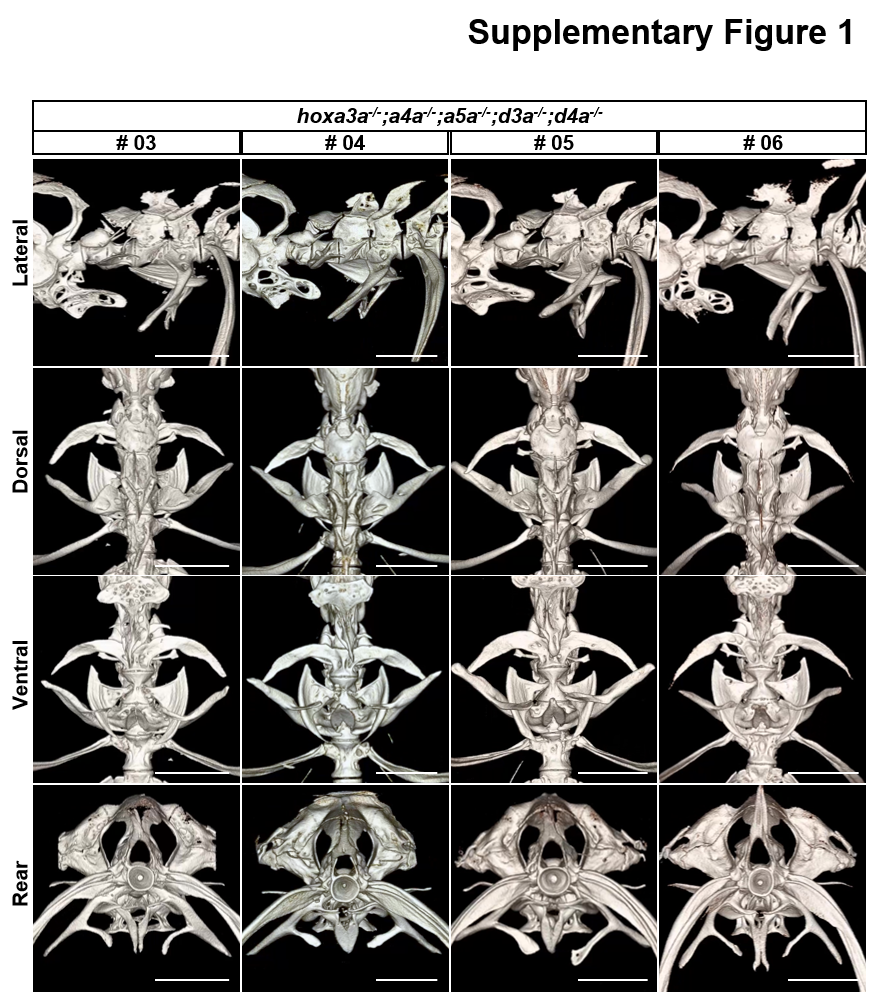

**Supplementary Figure S1. Anterior vertebral morphology in the remaining four *hoxa3a;a4a;a5a;d3a;d4a* homozygous mutants.**

X-ray micro-CT reconstructions of the anterior vertebrae and associated Weberian apparatus of mutant specimens #3–6, shown at orientations and magnifications comparable to those in Fig. 3. No obvious anterior vertebral transformations were detected in any of these four specimens. Scale bar: 1 mm.

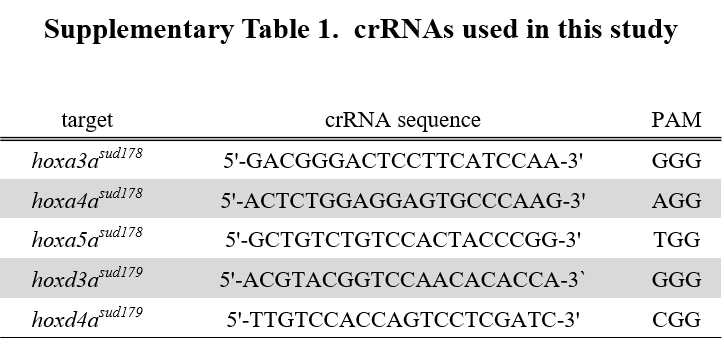

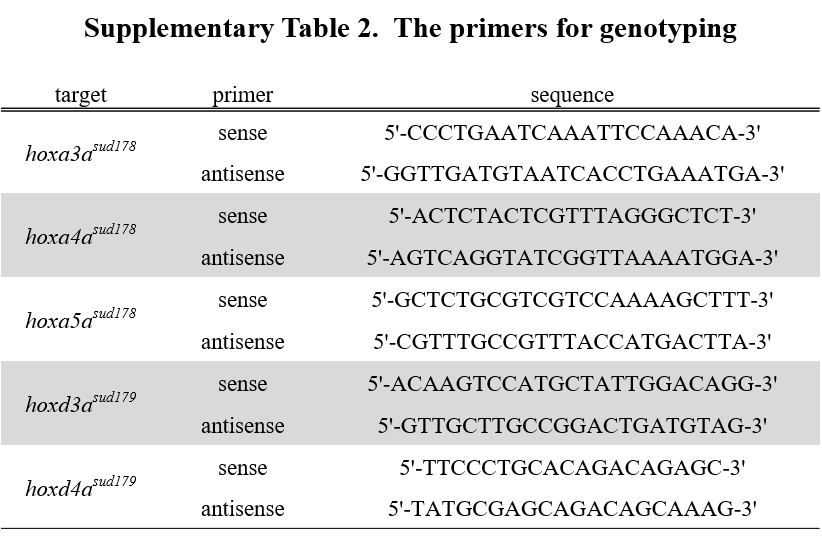

**Supplementary Movie 1. Three-dimensional reconstruction of the anterior vertebral column of wild-type zebrafish.**

Three-dimensional reconstruction generated from X-ray micro-CT data showing the anterior vertebrae and associated Weberian apparatus of an adult wild-type zebrafish.

**Supplementary Movie 2. Three-dimensional reconstruction of the anterior vertebral column of a *hoxaa*^−/−^; *hoxab*^+/−^; *hoxda*^+/−^ zebrafish.**

Three-dimensional reconstruction generated from X-ray micro-CT data showing the anterior vertebrae and associated Weberian apparatus of an adult *hoxaa^−/−^; hoxab^+/−^; hoxda^+/−^* zebrafish. No obvious alterations in anterior vertebral morphology are evident.

**Supplementary Movie 3. Three-dimensional reconstruction of the anterior vertebral column of a *hoxaa*^+/−^; *hoxab*^+/−^; *hoxda*^−/−^ zebrafish.**

Three-dimensional reconstruction generated from X-ray micro-CT data showing the anterior vertebrae and associated Weberian apparatus of an adult *hoxaa*^+/−^; *hoxab*^+/−^; *hoxda*^−/−^ zebrafish. No obvious alterations in anterior vertebral morphology are evident.

**Supplementary Movie 4. Three-dimensional reconstruction of the anterior vertebral column of a *hoxa3a;a4a;a5a;d3a;d4a* homozygous mutant.**

Three-dimensional reconstruction generated from X-ray micro-CT data showing the anterior vertebrae and associated Weberian apparatus of an adult *hoxa3a;a4a;a5a;d3a;d4a* homozygous mutant (#1). No obvious alterations in anterior vertebral morphology or Weberian ossicle identity were detected.
